# Spt5’s Central KOW Domains and the Pol II Stalk Collaborate to Regulate Chromatin and 3’-End Processing

**DOI:** 10.64898/2026.03.09.710576

**Authors:** Z. A. Morton, M. J. Doody, N. Naik, N. Paniagua, C. Delahunty, J. Yates, C.J. Bustamante, G. A. Hartzog

## Abstract

Spt5 is a universally conserved multidomain transcription elongation factor that acts as a component of all Pol II elongation complexes. Structural studies indicate that several of Spt5’s central KOW domains lie adjacent to the Pol II stalk, composed of subunits Rpb4 and Rpb7. However, their *in vivo* functions are unknown. Here we show that Spt5 and Rpb4/7 jointly modulate 3’-end formation and co-transcriptional chromatin integrity in *Saccharomyces cerevisiae*. We identify mutations in the *SPT5 KOW2-3* domains and *RPB7* that cause cryptic initiation of transcription and alter 3’-end formation of RNA transcripts. Molecular readthrough assays reveal allele-specific changes at both *GAL10* and *SNR13,* consistent with impacts on CPF/CF- and NNS-dependent termination. Proteomic experiments with isolated KOW2-3 domain enrich factors from both pathways as well as chromatin regulators, overlapping known Rpb7 interactors. Together, these findings support a model in which Spt5 KOW2-3/Pol II stalk region acts as a recruitment platform that coordinates pre-mRNA processing and chromatin dynamics during elongation, revealing new roles for the central KOW domains of Spt5.

**Summary:** This work describes a cooperative *in vivo* function for Spt5’s central KOW domains and the Pol II stalk in *Saccharomyces cerevisiae*. Allele-specific genetics and reporter assays show cooperative effects of *SPT5* and *RPB4/7* on cryptic initiation and 3′-end formation; double-mutant analyses reveal synthetic interactions. RT-qPCR at *GAL10* and *SNR13* demonstrates regulation of both poly(A) and non-coding transcript termination. Spt5 KOW pull-down proteomics enrich poly(A) and non-coding termination factors, as well as chromatin regulators that overlap with known Rpb7 interactors. Together, the data support a model in which Spt5 and the Pol II stalk coordinate chromatin integrity and termination during elongation.

## Introduction

Spt5 is an essential, multidomain transcription elongation factor conserved across bacteria, archaea, and eukaryotes. In archaea and eukaryotes, Spt5 has a conserved binding partner in Spt4, a small zinc-finger protein that stabilizes Spt5 (Malone et al. 1993; Ding et al. 2010). The bacterial homolog of Spt5, NusG, is composed of an N-terminal NGN domain and a C-terminal KOW domain. NusG’s NGN domain promotes transcription processivity by sealing DNA in the RNA polymerase central cleft and clamping RNA polymerase on DNA (Wang and Artsimovitch 2020). This function is conserved in eukaryotes. While NusG contains only a single KOW domain, eukaryotes have up to 7 (5 in *S. cerevisiae,* 4 to 7 in metazoans) (Pandey et al. 2023). While the conserved role of the NGN domain has been well-defined, the precise functions of Spt5’s central KOW domains remain ambiguous. As the expansion of Spt5’s KOW domains is eukaryote specific, we hypothesize that their function may be related to eukaryote specific transcriptional challenges, such as navigating chromatin or pre-mRNA processing. Indeed, in yeast, Spt5 is known to facilitate splicing, mRNA export, transcription through chromatin and chromatin maintenance (Lindstrom et al. 2003; Mayer et al. 2012; Hartzog and Fu 2013; Crickard et al. 2017; Evrin et al. 2022; Song and Chen 2022). Supporting this model, structural studies have placed several KOW domains adjacent to the Pol II stalk near the mRNA exit channel, far from the catalytic core of Pol II (Bernecky et al. 2017; Ehara et al. 2017).

Rpb4 and Rpb7 are core Pol II subunits that comprise the stalk region of Pol II in eukaryotes. These Pol II subunits are unique in that they can reversibly dissociate from the body of Pol II and are sub-stoichiometric to the remainder of Pol II (Tan et al. 2003). Rpb4/7 are involved in every step in transcription, from initiation to termination (Kolodziej et al. 1990; Edwards et al. 1991; Choder and Young 1993; Choder 2004; Sharma and Kumari 2013; Allepuz-Fuster et al. 2019; Calvo 2020), and have been shown to have roles in recruitment of 3’-end processing factors and polyadenylation site choice (Runner et al. 2008; Moqtaderi et al. 2026). Additionally, Rpb7 has been shown to bind nascent mRNA (Meka et al. 2005; Ujvári and Luse 2006). Structurally, the Pol II stalk protrudes from the foot domain of Pol II, and Rpb7 serves to lock Pol II into a more processive conformation (Armache et al. 2005). The exposed position of Rpb4/7 away from the body of Pol II suggests it can attract diverse functional units to the transcription complex (Runner et al. 2008; Sharma and Kumari 2013; Schier and Taatjes 2020). Because KOW2-4 lie alongside Rpb4/7 near the mRNA exit channel (Bernecky et al. 2017; Ehara et al. 2017; Qiu and Gilmour 2017), the combined KOW-stalk surface is well placed to impact termination pathway selection and mRNP maturation.

Here we test the hypothesis that the Spt5 KOW2-4/Pol II stalk region modulates co-transcriptional chromatin structure and 3’-end formation in yeast. We combine allele-specific genetics, molecular readthrough assays at *GAL10* and *SNR13,* and KOW-domain pulldown proteomics. We find that *spt5* KOW2-3 and *rpb7* alleles differentially affect cryptic initiation and polyadenylation (poly(A)) site choice as well as modulate CPF/CF (Cleavage and Polyadenylation Factor/Cleavage Factor) and NNS (Nrd1-Nab3-Sen1) mediated termination. Further, Spt5 KOW2-3 and KOW4 proteomics reveal enrichment for chromatin-related factors as well as factors involved in both canonical yeast termination pathways (CPF/CF and NNS) which overlap with known Rpb7 interactors, supporting a model in which the KOW-stalk surface serves as a context-dependent recruitment platform during elongation.

## Materials and Methods

### Media and yeast genetic methods

Yeast media and genetic manipulations followed standard methods (Rose *et al.,* 1990). Complete strain genotypes are listed in Table S1 (isogenic to S288C; *GAL2+*). Strain GHY1555 was a gift of Craig Kaplan and FY603 was a gift of Fred Winston. All other strains are from the Hartzog lab collection.

GHY4006 and GHY4007 were generated via a marker switch of GHY3349 and GHY3348 respectively harboring CKB310. PCR amplification of pRS404 was performed using OGH473 and OGH474 (1x Mango Mix (Bioline), 30 cycles of 95 °C 30 s, 51 °C 1min, 72 °C 90 s). High efficiency transformation was performed and cells were plated onto –Trp to select for plasmids in which *TRP1* replaced *LEU2.* Transformants were then replica plated to –Trp –Leu to screen for colonies that failed to grow, indicating loss of the *LEU2* marker on CKB310.

### Plasmids

A detailed list of the plasmids used in this study is provided in supporting information Table S2. Plasmid pZM6 was generated by PCR mutagenesis. Plasmid CKB223 was amplified with either OGH1630+OGH1631 or OGH1629+OGH1632 (Phusion polymerase, 1x Phusion HF Buffer (Thermo Scientific), 0.1 mM dNPTs; 30 cycles 95 °C 2min, 62 °C for 1min, 70 °C for 30 s). Gel excision was performed on the products using Nucleospin Gel and PCR cleanup kit (Macherey-Nagel). 5 ml of each product was added to a new reaction under the same conditions lacking primers and cycled 5 times (95 °C 30 s, 65 °C 1min, 70 °C 45 s). OGH1631 and OGH1632 were then added and cycled 30x. CKB233 and the PCR product were individually digested with *Xho*I and *Eag*I and the digested PCR product was ligated into the CKB233 backbone. To integrate *spt5-E546K* and *spt5-G587D*, a high efficiency transformation was performed with plasmids pMD11 and pMD18 that were digested with *Eag*I and *Hind*III into GHY3246 and GHY2741 respectively. Transformants were recovered after 24 h at 30 °C on YPD, replica plated to 5-FOA for counter-selection (2 x 48 h, 30 °C), then to 5-FOA –Trp. Stable Trp- colonies were restreaked and verified by PCR/Sanger sequencing.

### Oligonucleotides

A detailed list of oligonucleotides used in this study is provided in supporting information Table S3. OGH1773, OGH1774, OGH1775, OGH1776 sequences were originally used in Lee *et al.,* 2020

### Hydroxylamine Mutagenesis Based Screens

Plasmids were mutagenized with hydroxylamine as described (Rose, Fink 1987). Briefly, 10 mg of plasmid DNA was incubated in 1M hydroxylamine for 20 h at 37 °C. Reactions were quenched with 10 mL 5M NaCl and 50 mL 1 mg/ml BSA and ethanol precipitated. Pellets were resuspended in TE, re-precipitated, pelleted, air-dried and resuspended in 100 mL TE. Mutagenized DNA was used directly for transformation of *S. cerevisiae* by standard high-efficiency methods (GHY1555 for *RPB7 gal10Δ56,* GHY3244 for *RPB7* cryptic initiation, GHY2741 for *SPT5* cryptic initiation). Transformants were plated onto SC -Leu to select for transformants that contained the mutant *SPT5* or *RPB7* plasmids. Each plate was passed over 5-FOA 2 times to counter-select against the wildtype *URA3* plasmid. These plates were then replica plated to selective media (standard YP media with 2% galactose (YPGal) for *gal10Δ55/56,* SC -His with 2% galactose for cryptic initiation) and allowed to grow at 30 °C for 2 days to assay for the phenotype of interest. Colonies that grew on the selective media were re-streaked on YPD and then replica plated again to selective media and allowed to grow at 30 °C for 2 days to confirm the phenotype. SC -HIS + 2% glucose and YPGal were used as controls for cryptic initiation to ensure there was no leaky *HIS3* expression under non-inducing conditions. Plasmids were rescued and transformed into DMSO competent DH5α *E. coli,* retransformed into *S. cerevisiae* to confirm the phenotype is plasmid-dependent, and then subjected to Sanger sequencing. Phenotypes were finally scored via serial dilution assay on selective media.

### Serial Dilution Assay

1 x 10^7^ cells from an overnight culture (30 °C, YPD) were resuspended in 1 mL of sterile water. 5 10-fold serial dilutions were spotted onto agar plates containing the indicated media and allowed to grow for 48 h at 30 °C (or as specified).

### Spt5-KOW Domain Affinity Chromatography

BL21-DE3 cells harboring plasmids pGH382 (K2K3-His6) and pGH378 (L2K4-His6) were inoculated from a single colony in 10 mL LB+Kan at 37 °C and grown overnight on a rotator. 1 mL of culture was used to inoculate 1.3 L of LB+Kan at 37 °C and grown to OD 0.7 on an orbital shaker. Cultures were then induced with IPTG to 0.5mM and shaken at 20 °C overnight, harvested and frozen in liquid nitrogen. Frozen cell pellet was grown to a fine powder with a mortar and pestle frozen with liquid nitrogen and scooped into screw-cap tubes and frozen again in liquid nitrogen. 8 vials of cells were allowed to thaw on ice and mixed each with 0.5mL of phosphate buffer (P-Buffer) (50 mM phosphate pH 7.8, 10% glycerol, 50 mM NaCl, 50 mM Arginine, 50 mM Glutamate, PMSF 1x added fresh). Cells were then subjected to 3 rounds of sonication at 4 °C, each round being 30 s followed by 30 s on ice. Cells were spun for 10 min at 10,000 rpm in a microfuge at 4 °C. 1 mL HisPur Ni-NTA beads (Thermo Scientific) were equilibrated in P-Buffer, then placed in a chromatography column along with 4 mL of cell extract and placed on a rocker at 4 °C for 30 minutes. Column was washed with 20 CV P-Buffer at 250 mM NaCl, followed by 20 CV P-Buffer with 0.1% NP-40, followed by 20 CV P-Buffer with 100 mM NaCl and 4 mM imidazole. Column was eluted with 10 500 ml fractions of P-Buffer with 200 mM imidazole and frozen in liquid nitrogen. Fractions were then analyzed via SDS-PAGE. 5 Fractions containing the highest quantity of protein as determined by SDS-PAGE were buffer-exchanged and concentrated using Amicon Ultra-0.5 kDa MWCO centrifugal filters (3 exchanges; 4,000 rpm, 40 min each) into 25 mM MES. Purified protein (either K2K3, L2K4 or BSA) was incubated with Affi-Gel 10 resin with end-over-end mixing at 4 °C until the supernatant protein plateaued by Bradford (∼2.5 h). Reaction was quenched with 1M ethanolamine for 1 h at 4 °C. 1 mL of Affi-gel from each of K2K3, L2K4, or BSA couplings were transferred to separate chromatography columns, drained and washed with 10 CV of yeast lysis buffer. 25 mg crude yeast extract generated from GHY610 was diluted in 50 mL lysis buffer and loaded onto each column over a period of 8 hours. Columns were washed with 10 CV yeast lysis buffer with 0.5% NP-40 at 0.2 M K(Ac), then 5 CV yeast lysis buffer without NP-40. 8 400 ml fractions were eluted with a K(Ac) step-gradient (0.3-1.0 M in lysis buffer), starting with 0.3M and ending in 1M, followed by a 2.5 M urea strip. Fractions were split into 2 aliquots, frozen in liquid nitrogen and stored at −80 °C. Half of each fraction was TCA precipitated and analyzed via SDS-PAGE on a silver-stained gradient gel. Fractions containing 0.8M, 0.9M and 1M K(Ac) containing ∼8-10ug of protein were TCA precipitated and subject to MudPIT mass spectrometry analysis.

### Yeast Extract Preparation

Mid-log phase cells were harvested via centrifugation and frozen in liquid nitrogen. Cell pellets were ground into a fine powder using a mortar and pestle under liquid nitrogen. Cell powder was allowed to thaw on ice and then mixed with an equal volume of yeast lysis buffer (For affinity chromatography: 30 mM HEPES, pH 7.4, 200 mM potassium acetate (K(Ac)), 1 mM magnesium acetate, 1 mM EGTA, 0.05% Tween-20, 10% glycerol, 1 mM PMSF, 0.02 mM Pepstatin A, 0.02 mg/mL Chymostatin, 6 nM Leupeptin, 0.02 mM Benzamidine HCl, added fresh. For co-IP: 6 mM Na_2_PO_4_, 4 mM NaH_2_PO_4_·H_2_O, 1% NP-40, 150 mM NaCl, 2 mM EDTA, 1 mM EGTA, 50 mM NaF, 4 ug/mL Leupeptin, 0.1 mM Na_2_VO_4_, 5% glycerol. To 50 mL of buffer, 1 complete protease inhibitor cocktail tablet (Roche) was added along with 130 mL 0.5 M benzamidine, 500 mL 100 mM PMSF). Cells were then centrifuged at 14,000rpm for 10 minutes at 4 °C and supernatant collected. For affinity chromatography experiments, supernatant was clarified using an ultracentrifuge at 108,628 x g for 30 minutes at 4 °C.

### Mass Spectrometry

Sample pellets were resuspended in 60 uL of buffer (8 M urea 100 mM Tris pH 8.5) and reduced with 3 uL of 100 mM tris (2-carboxyethyl)phosphine hydrochloride (TCEP)). Samples were alkylated in the dark for 20 min with 250 mM iodoacetamide and digested with trypsin overnight at 37 C. Samples were acidified with formic acid (5%), and approximately 0.5 ug of sample was loaded onto EvoTips (Evosep) according to the manufacturer’s protocol. Samples were run on an Evosep One (Evosep) coupled to a timsTOF Pro mass spectrometer (Bruker Daltonics). Peptides were separated with a gradient of buffer A (0.1% formic acid in H2O) and buffer B (0.1% formic acid in acetonitrile) on a 15 cm × 150 μm ID column with BEH 1.7 μm C18 beads (Waters) and an integrated tip pulled inhouse. MS scans were acquired in PASEF mode, with 1 MS1 TIMS-MS survey scan and 10 PASEF MS/MS scans per 1.1 s acquisition cycle. Both ion accumulation time and ramp time in the dual TIMS analyzer were set to 100 ms, and the ion mobility range was 1/K0 = 0.6 to 1.6 V s cm−2. The m/z range was 100−1700. Precursor ions selected for MS/MS analysis were isolated with a 2 Th window for m/z < 700 and 3 Th window for m/z > 700. Collisional energy was lowered linearly from 59 eV at 1/K0 = 1.6 V s cm−2 to 20 eV at 1/K0 = 0.6 V s cm−2 as a function of increasing mobility. Precursors for MS/MS were picked at an intensity threshold of 2500, a target value of 20 000, and an active exclusion of 24s. Singly charged precursor ions were excluded with a polygon filter. Tandem mass spectra were extracted from raw files using RawExtract (Version 1.9.9) and were searched using ProLuCID against a standard UniProt *Saccharomyces cerevisiae* database concatenated with with reversed sequences. A static modification of carbamidomethylation on cysteine (57.02146) was considered. Data were searched with 50 ppm precursor ion tolerance and 600 ppm fragment ion tolerance. Data were filtered using DTASelect2 to a protein false-positive rate of < 1%. A minimum of 2 peptides per protein and 1 tryptic end per peptide were required. Statistical models for peptide mass modification (modstat) and tryptic status (trypstat) were applied.

### RNA Extraction and RT-qPCR Analysis

Cells were grown to mid-logarithmic phase in YPD and 10 mL of cells were harvested for RNA extraction. For analysis of *GAL10* transcripts, cells were grown in YP Raffinose to mid-logarithmic phase and induced with 2% galactose for 1 h before harvest. RNA was extracted with phenol:chloroform:isoamyl alcohol (25:24:1) (PCI) at 65 °C for 10 minutes with intermittent vortexing. RNA was purified, precipitated, and resuspended in RNase free water. Genomic DNA was removed using DNase I (New England Biolabs) and RNA was PCI purified, precipitated and resuspended in RNase free water. cDNA was prepared using combined random hexamer and oligo-dT primers (LunaScript RT SuperMix Kit, New England Biolabs) according to manufacturer instructions. qPCR amplification of cDNA was performed using Luna universal qPCR master mix according to manufacturer instructions using a Bio-Rad CFX96 Touch Real-Time PCR Detection System with melt-curve verification. Relative quantification used the DDCt method. All primer sequences are detailed in Supplementary Table 3. For *GAL10,* qPCR signal for the readthrough product (OGH1664, OGH1665) was normalized to the *GAL10* ORF (OGH342, OGH343). For *SNR13* (OGH1673, OGH1674), qPCR signal was normalized to *ACT1* (OGH1675, OGH1676). 3 biological replicates were assessed for each genotype. For *GAL10* readthrough experiments, strains used were GH1555, GHY3244, and GHY3368. For *SNR13* readthrough experiments, GHY3244, GHY3350, GHY4006 GHY4007, and GHY3368 were used. The indicated alleles of *RPB7* were transformed and the strain was passed over 5-FOA to counter-select for loss of the *RPB7 URA3* plasmid prior to analysis.

### Nrd1 Co-IP

2 mg of protein from crude yeast extract from FY603 (mock), GHY3244 + GHB464 (Rpb3-TAP), GHY3368 + GHB1417 (Rpb3-TAP, Spt5-E546K + Rpb7-D166G) were loaded onto 20 mL settled IgG Sepharose beads (Cytiva), pre-equilibrated in lysis buffer and end-over-end rotated for 2 h at 4 °C. Beads were washed 3 times with TEV wash buffer (10 mM Tris-HCl, pH 8.0, 150 mM NaCl, 0.1% NP-40) and resuspended in 40 mL of TEV wash buffer with 1 mM DTT. On-bead TEV cleavage was performed by adding 2 mL of in-house purified TEV protease (200 mM stock) followed by end-over-end rotation overnight at 4°C. Supernatant was removed and proteins were separated via 10% SDS-PAGE gel (acrylamide/bis) and transferred to nitrocellulose via wet transfer (100 V, 90 min, 4°C). Membrane was blocked with 5% milk in TBS-T for 1 hour, probed with rabbit anti-Nrd1 (1:5000) followed by goat anti-rabbit HRP (1:5000), stripped (.2M glycine, 0.1% SDS w/v, 1% Tween-20 v/v, pH 2.2) at RT 2 x 15 min, washed, re-blocked, and probed with goat anti-Rpb1 (1:2000) followed by donkey anti-goat HRP (1:5000). Images were collected using Bio-Rad Chemi Doc MP and analyzed using FIJI.

### Antibodies

Nrd1 antibody was generously provided by Dr. David Brow. Anti-Rabbit was acquired from Invitrogen (31460). Anti-goat antibody was acquired from Invitrogen (A15999). Rpb1 antibody was acquired from Santa Cruz Biotechnology.

## Results

### Spt5 Cryptic Initiation alleles map adjacent to the Pol II stalk and suppress *gal10Δ56*

We screened for *spt5* mutations that disrupt chromatin structure by utilizing the well-established *pGAL1-FLO8::HIS3* cryptic initiation reporter in which disruption of chromatin reveals an intragenic transcription start site that allows for growth on -HIS media (Cheung et al. 2008) (Fig. 1b). This screen yielded 3 mutations that alter residues within the KOW2-3 region of Spt5: G587D, G602S/S809F, and E546K (Fig. 1a). Spt5-G587 and -G602 pack closely at the juncture of Rpb7 and KOW3, while Spt5-E546 lies at the tip of KOW2 (Fig. 1d). Further characterization of these mutants and others identified from this screen will be described elsewhere.

**Figure 1:**
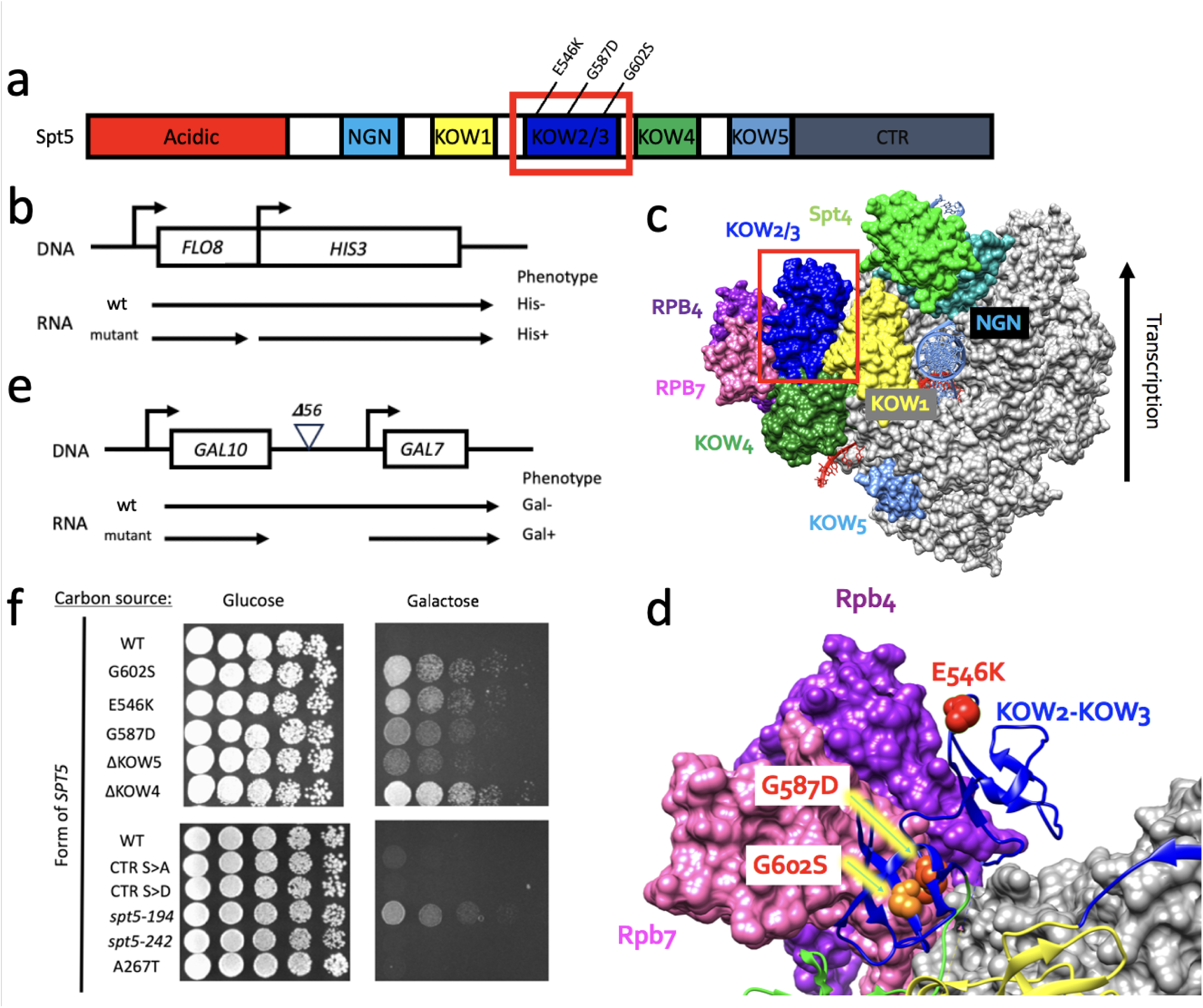
*spt5* CI Mutants Lie Adjacent to the Pol II Stalk and Suppress *gal10Δ56.* A) Spt5 domain architecture with cryptic initiation mutations indicated. B) Schematic of cryptic initiation reporter. Growth on –His media indicates cryptic initiation. C) cryo-EM structure (PDB: 5OIK) of Pol II EC reconstructed from *B. taurus* Pol II and *H. sapiens* Spt4/5, with KOW2-3 region boxed in red. Grey = Pol II, light green = Spt4, teal = Spt5-NGN, yellow = Spt5-KOW1, blue = KOW2/3, green = KOW4, cornflower blue = KOW5, pink = Rpb7, purple = Rpb4. Image generated in ChimeraX. (Bernecky et al. 2017) D) Zoomed in image of red boxed portion of panel C, indicating amino acid changes of cryptic initiation mutants (PDB: 5OIK) (Bernecky et al. 2017). Image generated in ChimeraX. E) Schematic of *gal10**Δ**56* reporter. Growth on galactose media indicates suppression of *gal10D56.* F) Serial 5-fold dilutions of a haploid yeast *spt5**Δ*** shuffle strain (GHY3246) with *gal10**Δ**56* containing the indicated alleles of *SPT5* on a centromere plasmid were spotted on YPD and YPGal plates and incubated at 30 °C.

Structural and biochemical studies of Pol II elongation complexes (ECs) show Spt5 KOW domains 2-4 lie adjacent to and contact Rpb4/7 (Fig. 1c) (Li et al. 2014; Bernecky et al. 2017; Ehara et al. 2017; Gong and Li 2023). We therefore hypothesized that the function of Spt5’s central KOW domains may be related to the known functions of Rpb4/7. Since loss of *RPB4* results in alteration of polyadenylation site choice, we wondered if our KOW2/3 mutations would also alter polyadenylation site choice (Runner et al. 2008; Moqtaderi et al. 2026). To test this, we utilized the *gal10****Δ****56* reporter gene (Kaplan et al. 2005) (Fig. 1e). This reporter truncates the *GAL10* poly(A) site, resulting in transcriptional readthrough into and interference of the *GAL7* promoter and yielding poor growth on galactose. Mutations that perturb polyadenylation result in termination at the truncated poly(A) site and restore growth on galactose media. All 3 point mutations in KOW2-3 resulted in suppression of *gal10****Δ****56*, as did a deletion of KOW4 and, to a lesser degree, KOW5 (Fig. 1f). To confirm that this functionality is specific to the KOW domains we also tested previously described mutations in the N- and C-terminal regions of Spt5. With the exception of *spt5-194*, which has previously been shown to destabilize Spt5’s overall structure (Ding et al. 2010), these *spt5* mutations did not suppress *gal10****Δ****56*.

### Mutations in *rpb4/7* result in altered poly(A) site choice, cryptic initiation, and genetically interact with *spt5-KOW* mutants

We next asked if *rpb4/7* mutations share phenotypes with *spt5-KOW2/3* mutations. As *RPB7* is an essential gene, we mutagenized plasmid-borne *RPB7* and screened for cryptic initiation and suppression of *gal10****Δ****55,* which is similar to *gal10****Δ****56* (Greger and Proudfoot 1998; Kaplan et al. 2005). 2 mutations resulted in strong suppression of *gal10****Δ****55* – *rpb7-G149*D, and *rpb7-E100K* (Fig. 2b). Rpb7-G149 lies at the juncture of Rpb7 and KOW3 and sits near Spt5-G587, while Rpb7-E100 lies at the interface of Rpb7 and Spt5-KOW4 (Fig. 2d). We did not identify *rpb7* alleles that caused cryptic initiation in this screen. However, we also tested *rpb7* alleles generously provided by Craig Kaplan, which have been shown to exhibit transcriptional defects (*rpb7-D166G, rpb7-L168S, rpb7-V101E)* for suppression of *gal10****Δ****55* as well as cryptic initiation (Braberg et al. 2013). One of these alleles, *rpb7-D166G*, demonstrated strong cryptic initiation (Fig. 2b). Similar to Spt5-E546, Rpb7-D166 is a solvent-exposed residue that projects away from the polymerase (Fig. 2d). *Rpb7-D166G* was also shown to result in transcription start site (TSS) defects as well as sensitivity to mycophenolic acid (MPA), suggestive of defects in transcription initiation or elongation (Braberg et al. 2013). Furthermore, *rpb7-L168S* exhibited weak cryptic initiation, while *rpb7-V101E* did not suppress *gal10****Δ****55* or show cryptic initiation. This difference suggests that the cryptic initiation phenotypes seen in *rpb7-D166G* and *-L168S* are allele-specific, rather than resulting from general loss of function. Since Rpb7-D166G abolishes the negative charge of aspartate, we reasoned that a charge reversal may result in a stronger phenotype. We therefore engineered *rpb7-D166K* via PCR mutagenesis. However, contrary to our expectations, we found this allele did not give any observable phenotype as a single mutant.

**Figure 2:**
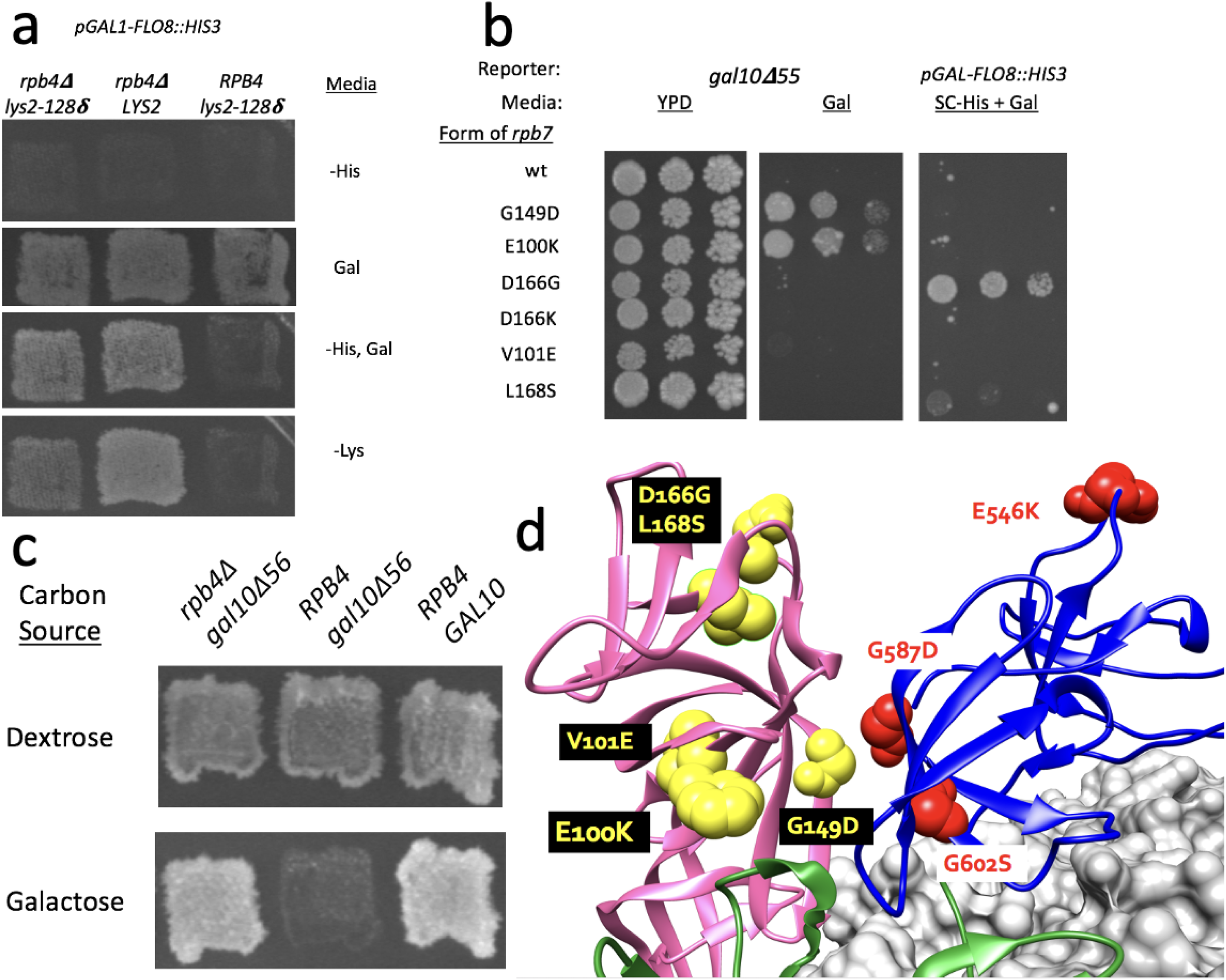
Mutations in *rpb4/7* result in cryptic initiation and suppression of *gal10Δ56.* A) Patches of haploid yeast strains (GHY4009, GHY4008, and GHY1934) harboring the cryptic initiation reporter gene construct and the indicated alleles of *RPB4* and *LYS2,* grown on the indicated media at 30 °C. Growth of *lys2-128δ* strains on–Lys media indicates an Spt- phenotype. B) Serial 5-fold dilutions of haploid yeast *rpb7* shuffle strains containing either *gal10**Δ**55* (GHY1555) or the cryptic initiation reporter (GHY3244) and harboring the indicated alleles of *RPB7* on a centromere plasmid were spotted onto the indicated media and incubated at 30°C. C) Patches of haploid yeast strains harboring the indicated alleles of *RPB4* and *GAL10* (GHY4005, GHY3147, GHY1934) were grown on the indicated media at 30 °C. D) Cryo-EM representation of *spt5* and *rpb7* alleles mapped to the Pol II EC. Spt5-KOW2/3 (blue), Spt5-KOW4 (green), Rpb7 (pink) and Pol II (grey) are shown (PDB: 5OIK) (Bernecky et al. 2017).

To assess the evolutionary and functional significance of these Spt5 and Rpb7 residues, we generated sequence alignments across model organisms and asked if these residues were highly conserved (Supplemental Fig. 1). Spt5-G602 is universally conserved as are the closely spaced Spt5-G587 and Rpb7-G149 residues. This suggests that this structure is evolutionarily relevant. Both Spt5-E546 and Rpb7-D166 appear only as either glutamate or aspartate, indicating that preservation of the acidic nature of these residues is universal. Rpb7-L168 is also universally conserved, highlighting the evolutionary importance of this structure formed by the C-terminus of Rpb7 and Spt5-KOW2.

Since *RPB4* is not essential, we deleted *RPB4* in strains carrying the cryptic initiation and *gal10****Δ****56* reporters and observed both suppression of *gal10****Δ****56* and cryptic initiation (Fig. 2a, c).

To further assess the functional cooperation of Spt5 KOW2-3 with the Pol II stalk, we generated double mutants of *spt5-E546K* and *spt5-G587D* with our previously identified *rpb7* mutants. Since *spt5-G602S* is also paired with *spt5-S809F*, we excluded this mutant from analysis as it would confound mechanistic interpretations of our results. We first tested our *rpb7* alleles in combination with *spt5-G587D* in the cryptic initiation background. *Spt5-G587D*, which displays a modest cryptic initiation phenotype when plasmid-borne (data not shown), showed a slightly attenuated phenotype when integrated. When combined with *spt5-G587D*, we observed a strong synthetic cryptic initiation phenotype in *rpb7-G149D*, *-E100K*, and *-D166K* (Fig. 3a). Interestingly, the cryptic initiation phenotype observed in the *rpb7-D166G* single mutant (Fig. 2b) was partially suppressed in the presence of *spt5-G587D*. Additionally, we observed strong synthetic growth defects at 39 ° C when *spt5-G587D* was combined with *rpb7-G149D, -D166G and -V101E*.

**Figure 3:**
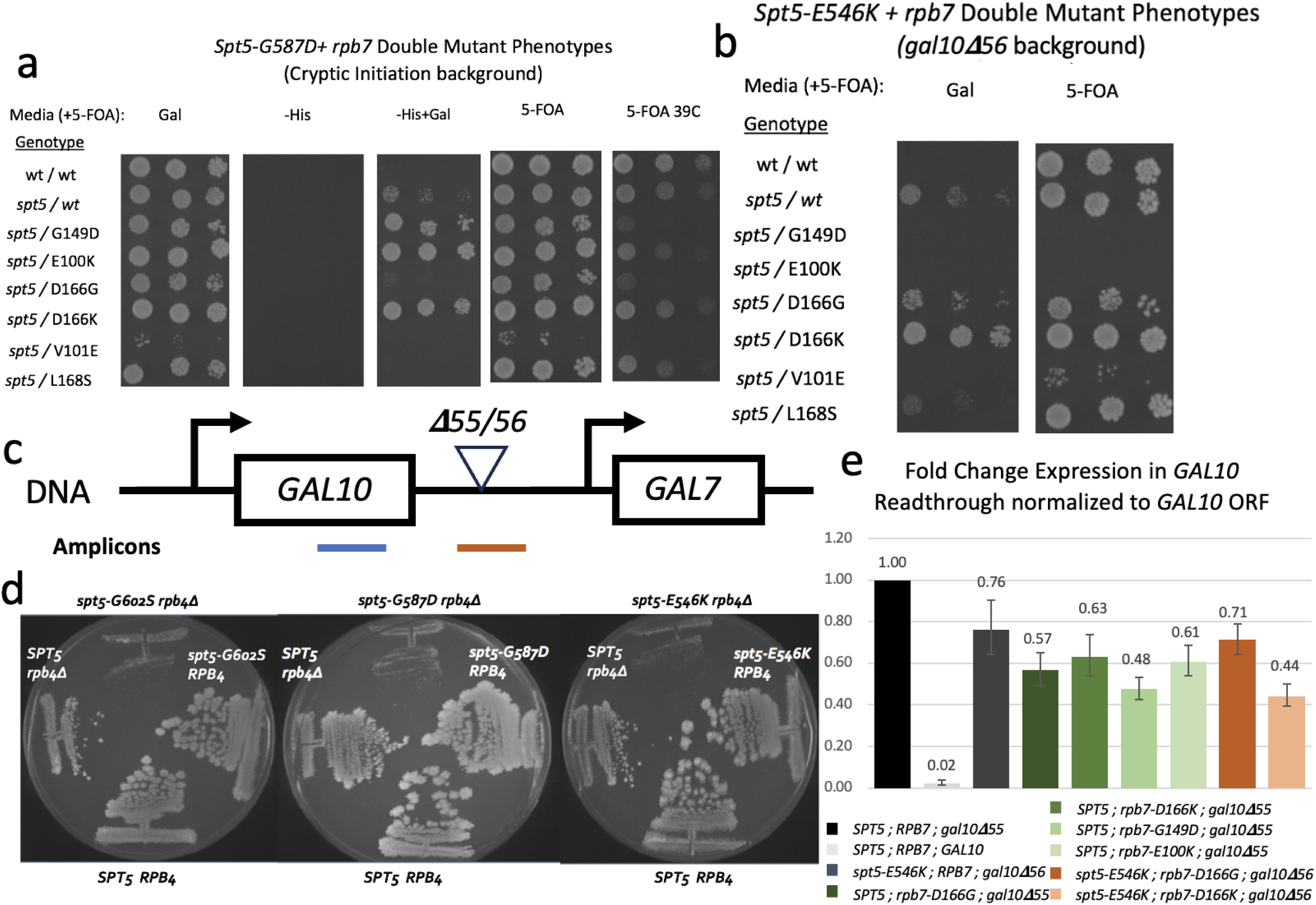
*SPT5* KOW2-3 and *RPB4/7* Genetically and Functionally Interact. A) Serial 5-fold dilutions of *rpb7 spt5-G587D* double mutants. A haploid *rpb7**Δ** spt5-G587D* strain with the cryptic initiation reporter and an *RPB7 URA3* plasmid (GHY3348) was transformed with centromeric plasmids carrying the indicated alleles of *RPB7* and passed over 5-FOA to select for loss of the *RPB7 URA3* plasmid. A haploid *rpb7**Δ** SPT5+* strain harboring a wildtype *RPB7 LEU2* plasmid was used as a control (GHY3243). These strains were spotted onto the indicated media and incubated at 30 °C B) Serial 5-fold dilutions of haploid yeast *rpb7**Δ*** shuffle strain with *spt5-E546K* and *gal10**Δ**56* (GHY3349) harboring the indicated alleles of *RPB7* on a centromere plasmid were spotted onto the indicated media and allowed to grow at 30 °C. A haploid *rpb7**Δ** SPT5 gal10**Δ**55* strain harboring a wildtype *RPB7 LEU2* plasmid was used as a control. *C*) Schematic detailing location of PCR amplicons on the *gal10**Δ**56/gal10Δ55* gene constructs. Blue amplicon is within *GAL10* ORF, red amplicon spans *GAL10* poly(A) site. D) Haploid *spt5**Δ*** shuffle strains harboring either wildtype *RPB4* (GHY2741) or *rpb4**Δ*** (GHY3248) and the indicated alleles of *SPT5* on a centromere plasmid were struck out on –Leu media and replica plated to 5-fluoroorotic acid (5-FOA) for counterselection against the *SPT5 URA3* plasmid. E) Bar chart indicating mean of n=3 biological replicate **ΔΔ**Ct values ± SD of *GAL10* readthrough product normalized to total *GAL10* transcripts relative to *SPT5; RPB7; gal10**Δ**55*. Significance between *gal10**Δ**55; SPT5; RPB7* and all other backgrounds was calculated by 2-tailed Student’s t-test; all p-values are < .002. Error bars indicate SD.

We next tested *rpb7* alleles in combination with s*pt5-E546K* in the *gal10****Δ****56* background (Fig. 3b). There was little if any additional effect when *spt5-E546K was* paired with *rpb7-D166G*, however the suppression of *gal10****Δ****56* was clearly enhanced when paired with *rpb7-D166K*. In contrast, suppression of *gal10****Δ****56* was reduced when *spt5-E546K* was combined with *rpb7-L168S*, and *spt5-E546K* resulted in inviability when combined with *rpb7-G149D* or-*E100K.* When we combined *spt5-G602S + S809F*, *spt5-E546K*, or *spt5-G587D* with a deletion of *RPB4*, we observed that all 3 double mutants were inviable (Fig. 3d).

### Molecular Readthrough at *GAL10* corroborates altered poly(A) site choice

To further investigate the genetic suppression of *gal10****Δ****56* observed in *spt5* and *rpb7* mutants, we utilized RT-qPCR to quantify readthrough of *GAL10* mRNA in *gal10****Δ****56* strains. We designed 2 sets of primers. The first falls within the *GAL10* ORF and will detect both readthrough and normally terminated *GAL10* transcripts. The second primer pair spans the *gal10****Δ****56* mutation and will only detect readthrough transcripts (Fig. 3c). To calculate the ratio of *GAL10* transcripts that fail to terminate properly, we calculated the ratio of the readthrough product to total *GAL10* mRNA normalized to the readthrough:total GAL10 ratio measured in *SPT5 RPB7 gal10****Δ****56* strain. A ratio < 1 indicates reduced readthrough. We found that *spt5-E546K* resulted in 0.76-fold readthrough relative to wild type*, rpb7-D166G* resulted in 0.57-fold readthrough, and -*D166K* resulted in 0.63-fold readthrough relative to wild type (Fig. 3e). We found that the *rpb7*-*G149D* and -*E100K* alleles resulted in a significant 0.48- and 0.61-fold reduction in readthrough, reflective of their strong suppression of *gal10****Δ****56.* We further found that when combined with *spt5-E546K, rpb7-D166G* resulted in 0.71-fold readthrough relative to wild type, which is consistent with the lack of additive effects seen in the genetic assay. *Spt5*-*E546K* + *Rpb7-D166K* showed significantly reduced readthrough that was lower than either single mutant alone at 0.44-fold, which also reflects the enhanced suppression in the genetic assay. Interestingly, while *rpb7-D166G* resulted in similar reduction in readthrough to other mutants, it did not suppress *gal10****Δ****55* on its own. Aside from this exception, the molecular results observed in our RT-qPCR assay are in close alignment with the genetic suppression observed with *gal10****Δ****55/56*.

### Spt5 KOW pulldown proteomics enrich factors from CPF/CF and NNS Termination Complexes as well as chromatin related factors

Proteomic and biochemical studies have identified a number of factors that interact with Spt5 (Krogan et al. 2002; Lindstrom et al. 2003). However, these binding events have not been assigned to individual Spt5 domains. Since Spt5’s domains are widely distributed across the surface of Pol II, it is reasonable to suspect that the different Spt5-interacting factors may bind exclusively to specific domains of Spt5 to support location-specific roles. Supporting this hypothesis, Spt5’s KOW domains are composed of Tudor folds which, in other proteins, are known to support protein-protein and protein-nucleic acid interactions (Lasko 2010; Meyer et al. 2015). To pinpoint the function of the conjoined Spt5 KOW-stalk region of the elongation complex, it is therefore necessary to identify factors that bind specifically to the KOW domains that participate in this structure.

To accomplish this, we performed affinity chromatography followed by MudPIT mass spectrometry to identify proteins captured by either purified KOW2-3 (K2K3) region, Linker2-KOW4 (L2K4) region, or BSA. To this end, we purified recombinant His-tagged K2K3 and L2K4 and BSA and immobilized them on Affi-Gel. The column was then challenged with a clarified yeast lysate, washed, and eluted via salt gradient. Silver-stained gels of the eluates revealed that the KOW2-3 and L2K4 bound noticeably different patterns of proteins (Supplemental Figure 2). The 3 highest salt fractions from K2K3, L2K4 and BSA were subjected to mass spectrometry.

We identified many proteins that were differentially enriched by KOW2-3 and L2K4, as well as proteins that bound both domains. We excluded a number of proteins from analysis: ribosomal proteins, which are frequently identified nonspecifically in mass spectrometry, cytoplasmic proteins, as well as importins and mitochondrial membrane proteins. Spt5 is known to interact with RNA Polymerase I, and we did identify a number of RNA Polymerase I subunits, however we excluded ribosome biogenesis factors as this study focuses on Pol II transcription. Finally, any proteins that were enriched less than two-fold relative to the BSA control column were excluded. These exclusions left 80 proteins that were enriched in either K2K3 or L2K4 (Table 1). This list includes proteins involved in a wide array of nuclear functions, including transcription, DNA replication, and splicing. Consistent with the cryptic initiation phenotype observed in our *rpb4/7* and *spt5* KOW2-3 mutants, we also identified proteins involved in chromatin structure and 3’-end formation, including Nap1, Nbp2, Histones H3, H2A.Z, H2B and Spt16. Interestingly, we also identified factors in both NNS and CPF/CF termination pathways. Prior proteomics studies and two-hybrid screens have identified factors that bind to Rpb7 (Mitsuzawa et al. 2003; Mosley et al. 2013). Many of these identified proteins overlap with factors identified in our affinity chromatography experiments, which are indicated in bold. Of particular interest is Nrd1, a component of the Nrd1-Nab3-Sen1 (NNS) complex which is involved in non-polyadenylated transcript termination, exosome-mediated RNA degradation and mRNA surveillance (Steinmetz and Brow 1996; Steinmetz and Brow 1998; Steinmetz et al. 2001; Vasiljeva and Buratowski 2006; Singh et al. 2021). Both Nrd1 and Nab3 were enriched by KOW2-3 in our study, and Nrd1 was identified in prior studies as interacting with Rpb7, indicating that the region comprised of Spt5 KOW2-3 and Rpb4/7 is a possible binding target for the Nrd1-Nab3-Sen1 termination complex (Mitsuzawa et al. 2003; Mosley et al. 2013).

**Table 1:**
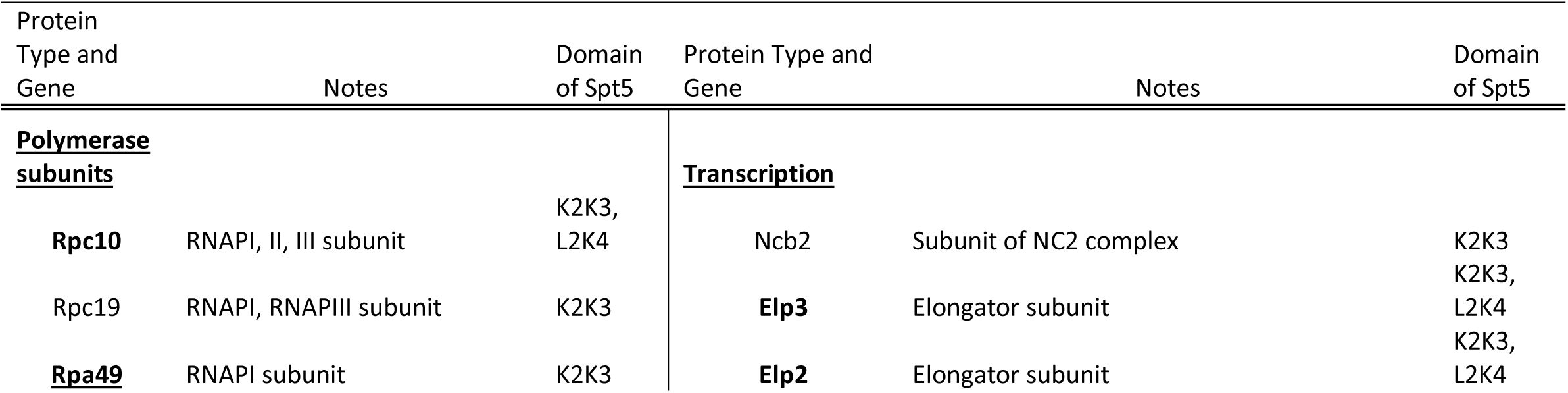
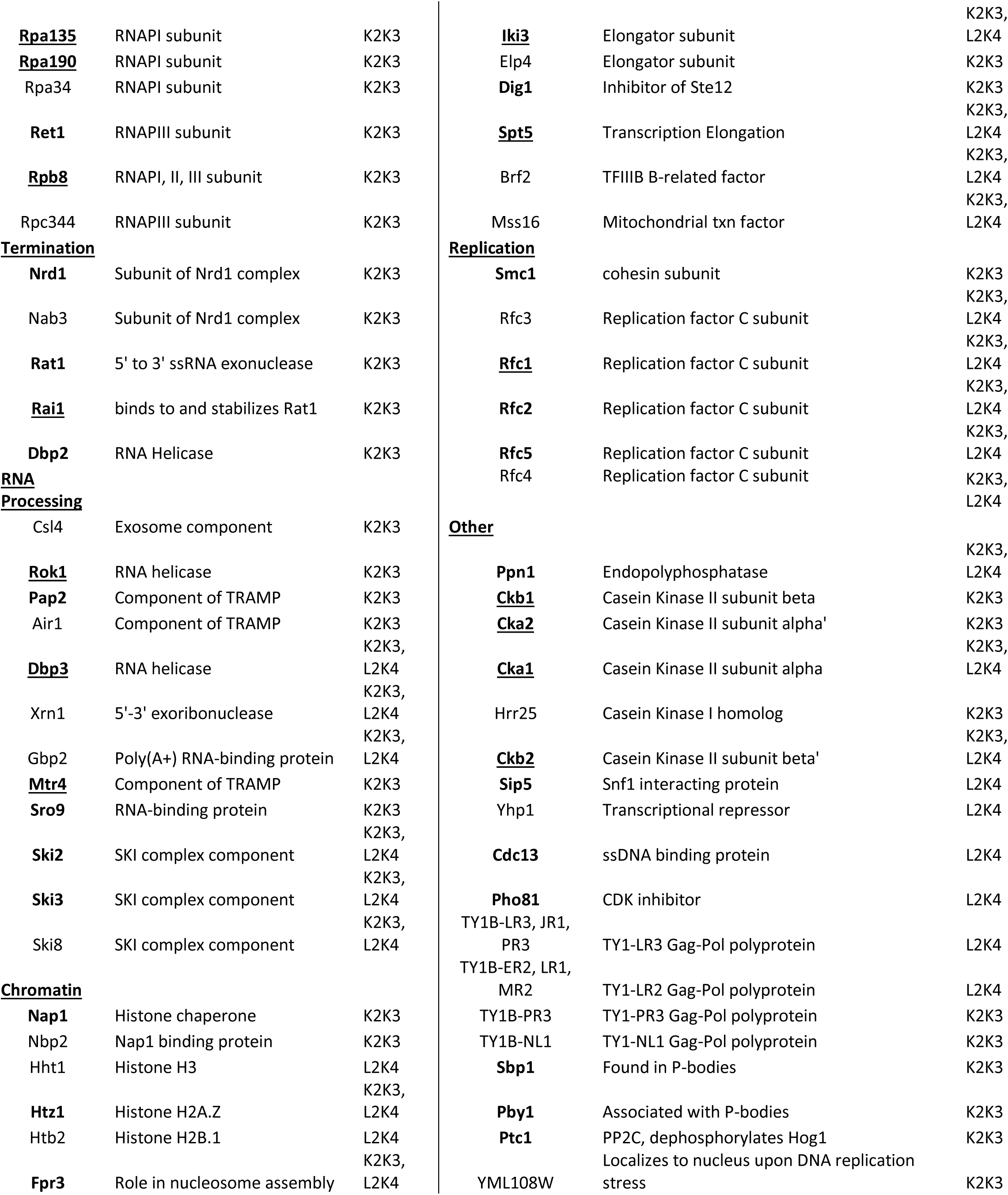
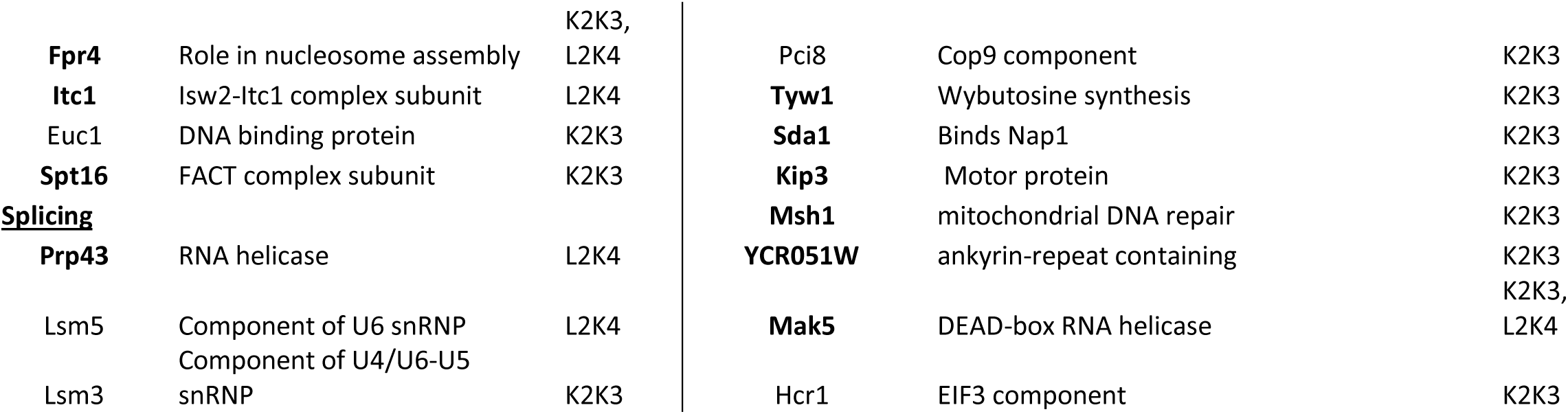
Identification of Proteins that Interact with Spt5’s Central KOW Domains. List of proteins identified via MudPIT mass spectrometry that are enriched at least 2-fold relative to BSA in pull-downs against the indicated KOW domain, excluding cytoplasmic proteins, ribosomal biogenesis factors, importins and mitochondrial proteins from the 1 M salt fraction. Proteins indicated in bold were previously identified as Rpb7-interacting (Mitsuzawa et al. 2003; Mosley et al. 2013) Underlined proteins were previously identified as Spt5-interacting (Lindstrom et al. 2003).

### Allele-specific regulation of NNS termination at *SNR13* by Spt5 and Rpb7

Given the enrichment of Nrd1 in the KOW2-3 pulldown, we asked if *NRD1* overexpression could suppress termination defects of our *spt5* mutations. We transformed yeast harboring the *gal10****Δ****56* reporter and *spt5* mutations with a *NRD1* overexpression vector and assayed for changes in phenotypes observed in the *spt5* single mutants. We observed modest suppression of both the *gal10****Δ****56* and the Spt- phenotype in *spt5-E546K,* indicating that Nrd1 may be targeting Spt5-E546 (Fig. 4a). Prior studies have shown that a *nrd1* mutation attenuates transcription termination at the *SNR13* (Lee et al. 2020). *SNR13* is a snoRNA gene that lies adjacent to *TRS31* and is regulated by NNS-dependent termination. When NNS-dependent termination is defective, transcription will read through into the adjacent *TRS31* gene resulting in a longer, bicistronic transcript. To test the model that the Spt5 KOW2-3/stalk region regulates NNS-dependent termination, we asked if mutations in *spt5* and *rpb7* would result in readthrough at *SNR13* into the downstream *TRS31.* To this end, we used RT-qPCR with a previously published primer pair that spans *SNR13* and *TRS31* to quantify total levels of the readthrough transcript, normalized to *ACT1* (Fig. 4b) (Lee et al. 2020). Consistent with our expectations, we found that *rpb7-D166G* resulted in two-fold enhanced readthrough relative to wild type (Fig. 4c). Further, we found that the *rpb7-D166G, spt5-E546K* double mutant further enhanced readthrough to roughly 5-fold relative to wild type. We also found that *spt5-E546K, spt5-G587D, rpb7-E100K* and *rpb7-G149D* resulted in decreased readthrough, indicating enhanced 3’-end formation, consistent with our observations in *gal10****Δ****56.* Interestingly, the *spt5-G587D, rpb7-D166G* double mutant resulted in suppression of readthrough to wild type levels, which is consistent with our observation that this double mutant resulted in suppression of cryptic initiation. To test if this double mutant disrupts recruitment of Nrd1 to the Pol II EC, we performed a pull-down experiment targeting TAP-tagged Rpb3 in wild type and mutant contexts probing for Nrd1 via Western blot. We found that there was no difference in the amount of Nrd1 associated with Pol II in the double mutant context compared to wild type, indicating that bulk Nrd1-Pol II association is retained (Fig. 4d).

**Figure 4:**
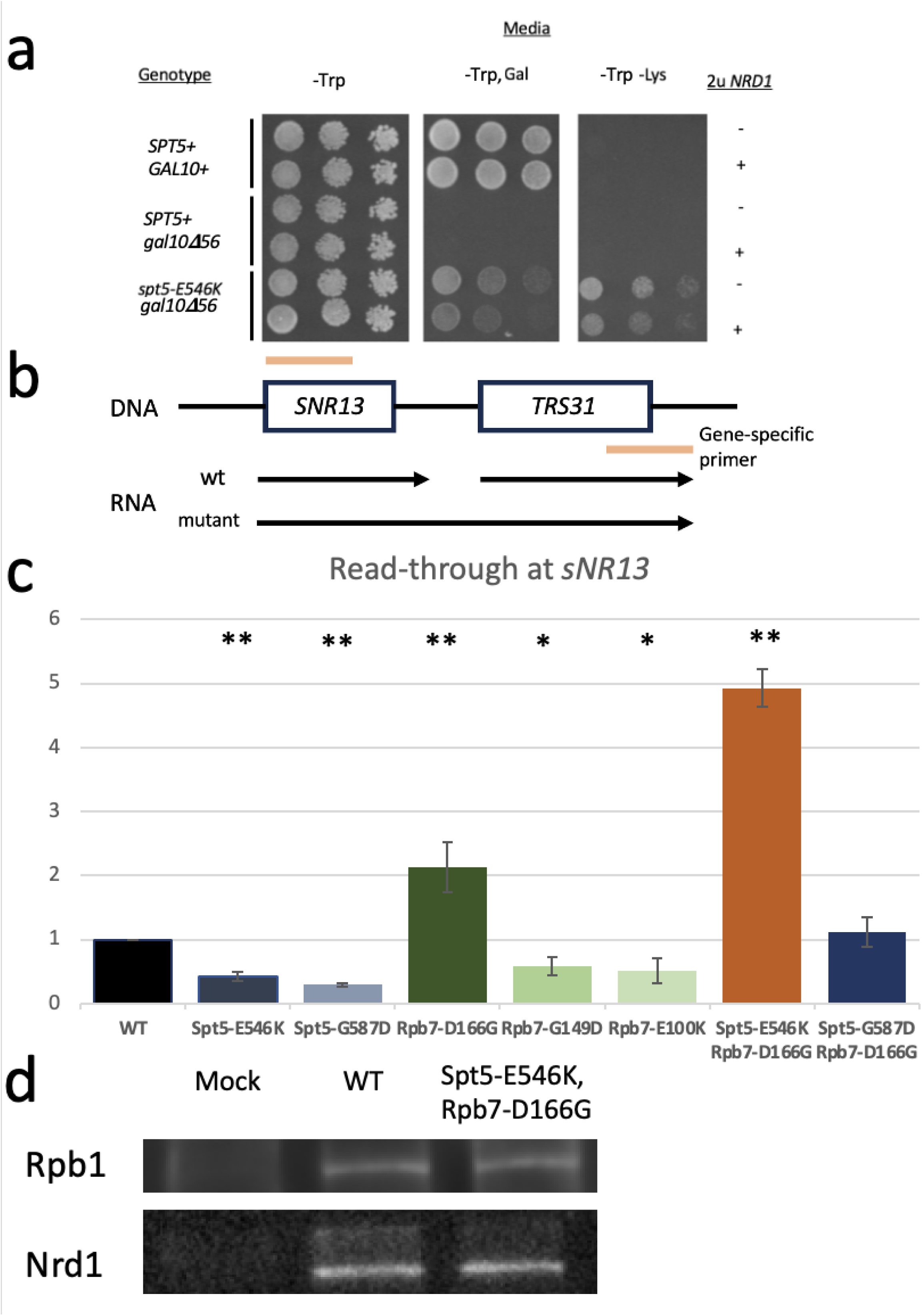
*spt5/rpb7* Mutants Alter non-coding RNA Termination and Interact Genetically with *NRD1* Without Disrupting Binding to Pol II. A) Serial 5-fold dilutions of haploid yeast strains (GHY1934, GHY3147, GHY3335) with the indicated allele of *GAL10* and *SPT5,* with and without *NRD1* supplied on a high-copy vector (2μ) were spotted onto the indicated media and incubated at 30 °C. B) Schematic showing locations of primers at the *SNR13, TRS31* locus. C) Bar chart showing mean of n=3 biological replicate **ΔΔ**Ct values ± SD of readthrough product at *SNR13* normalized to *ACT1* compared to wildtype. 2-tailed Student’s t-test was performed comparing WT to all other conditions to assess significance: * indicates p < 0.05, ** indicates p < 0.01. Error bars indicate SD. D) Western blot with equal inputs of Rpb3-TAP pull-down TEV-treated eluates in mock (untagged), WT and *spt5-E546K rpb7-D166G* genetic backgrounds, with primary antibody indicated on the left. Blots were stripped in between probing for Nrd1 and Rpb1. Representative of n=3 biological replicates. Strains used were FY603, GHY3244, and GHY3368. GHY3368 was transformed with CKB310 and passed over 5-FOA prior to analysis.

## Discussion

As Pol II traverses the length of a gene, the stalk region and its interacting factors are likely remodeled to better serve the elongation complex in response to co-transcriptional challenges, such as chromatin, mRNP packaging, and 3’-end processing (Mosley et al. 2013). Part of this remodeling procedure may involve a shift in conformation of Spt5’s central KOW domains or transient dissociation of Rpb4/7. Prior studies have shown that Pol II ECs lacking Rpb4/7 are enriched for specific elongation factors, and Pol II ECs purified via affinity-tagged Rpb7 are depleted for serine-2 CTD phosphorylation, suggesting that Rpb4/7 may dissociate in response to specific signals during elongation (Mosley et al. 2013). In fact, recent studies of native Pol II purified from *D. melanogaster* embryos have elucidated the structure of a stalk-less elongation complex actively engaged with and transcribing DNA, supporting the idea that 10-subunit Pol II has *in vivo* functions (Venette-Smith et al. 2025). Our genetic analysis of *spt5* and *rpb7* mutants supports a model in which the spatial arrangement of Rpb7 and Spt5’s central KOW domains relative to each other is functionally relevant. Importantly, the consequences of perturbing this surface are allele-specific: while Rpb7-G149 and Spt5-G587 are adjacent to each other according to many cryo-EM models, *rpb7-G149D* did not share the same phenotype as *spt5-G587D.* If this were the only relevant conformation, one would expect that mutating either *RPB7* or *SPT5* would result in the same phenotype. However, we observe that *rpb7-G149D* only resulted in a *gal10****Δ****55* phenotype, while *spt5-G587D* confers defects both in *gal10****Δ****56* and cryptic initiation. This suggests that merely breaking contact between Spt5 and Rpb7 is not sufficient to induce cryptic initiation, and that Spt5-KOW2-3 may have an independent function when not in contact with Rpb7. Despite this separation of function, our observation that *spt5-E546K* is inviable in the presence of *rpb7-E100K* or *rpb7-G149D* argues that the Spt5-stalk surface functions cooperatively, and disruption of this surface compromises an essential function. Whether the observed phenotypes are a result of induced dissociation of Rpb4/7 or conformational shifts of KOW2-4 will require future work. One possibility is that Spt5 contributes to retention or stabilization of the Pol II stalk during transcription.

Our finding that the *rpb7-D166G, spt5-E546K* double mutant does not impair recruitment of Nrd1 to Pol II is consistent with the existence of redundant recruitment mechanisms, including interactions with the Spt5 CTR and recognition of serine-5 phosphorylation of the Pol II CTD via the Nrd1 CID (Leporé and Lafontaine 2011; Heo et al. 2013). Although these mutations do not prevent NNS recruitment to Pol II, they may influence the productive engagement of NNS factors with the elongation complex or with nascent RNA. Previous work in *Schizosaccharomyces pombe* has shown that Rpb7-D167, which aligns with *S. cerevisiae* Rpb7-D166, is required for maintaining contact with Seb1, the *S. pombe* ortholog of Nrd1, and this interaction is conserved in *Saccharomyces cerevisiae* (Mitsuzawa et al. 2003). This suggests that the phenotypes we observe may result from altered positioning of NNS factors relative to the KOW-stalk surface.

Mutations in *SPT6* and other elongation factors have been shown to enhance upstream poly(A) site utilization at *GAL10* and other loci (Cui and Denis 2003; Kaplan et al. 2005; Geisberg et al. 2024; Moqtaderi et al. 2026). One model to explain this phenotype is that defects in transcription elongation result in a slower, more pause-prone polymerase, allowing for an increase in processing time at a weakened poly(A) site. Indeed, polymerase speed has been correlated with differences in poly(A) profiles across many genes, with a slower polymerase favoring upstream poly(A) sites, and a fast polymerase favoring downstream poly(A) sites (Geisberg et al. 2020; Geisberg et al. 2022). Further, defects in chromatin-related proteins such as Spt6 and FACT have been shown to shift poly(A) site choice upstream through effects on elongation rate arising from chromatin perturbations (Geisberg et al. 2024). However, these observations cannot completely explain the alterations in poly(A) site choice at *GAL10.* Kaplan *et al*. reported that in an *spt6* mutant the upstream poly(A) site is equally favored over 2 downstream alternative poly(A) sites (Kaplan et al. 2005). For this phenotype to be strictly explained by a slower elongating Pol II at *GAL10*, one would expect a gradient of decreasing poly(A) site usage following the initial Poly(A) site. This suggests that an alternative model may be true: suppression of *gal10****Δ****56* by perturbing elongation factors is caused by an alteration in either conformation or composition of the elongation complex. Supporting this, recent cryo-EM experiments have elucidated the structure of Pol II bound with the exonuclease Rat1, its binding partner Rai1, and Spt5 (Zeng et al. 2024). Interestingly, Pol II complexes that resolved Rat1 and Rai1 only showed density for Spt5-KOW5. In these structures, Rat1 binds Rpb7 at the same general location as Spt5 KOW2-3 and appears to contact Rpb7-G149, suggesting that Rat1/Rai1 trigger eviction of Spt5 KOW2-3 without disrupting general Spt5 association to Pol II. Single-molecule studies have also shown that Spt4/5 reversibly dissociates from the EC during a single transcription cycle (Rosen et al. 2020). These observations support the model that Spt5’s central KOW domains may indeed shift in conformation in response to the needs of the transcription complex. Taken together, these results suggest that the *spt5* and *rpb7* mutations may disrupt the interaction between Spt5 KOW2-3 and the stalk. Interestingly, *rpb7-G149D* was recently independently isolated in an unbiased screen for mutations that derepress transcription-coupled repair (TCR) (Gong and Li 2023). This group utilized photoreactive crosslinking to capture Rpb7-Spt5 interactions with wildtype Rpb7 and Rpb7-G149D. In the mutant context, they observed disruption of crosslinking between Rpb7-E148, Rpb7-I151 and Spt5. They also observed disruption of crosslinking between Rpb7-E100 and Spt5 in Rpb7-G149D, potentially explaining why *rpb7-G149D* and *rpb7-E100K* mirror each other’s phenotypes. Taken together, this supports the hypothesis that disrupting the contact between KOW2-3 and Rpb4/7 favors an alternate conformation of the Pol II EC that more strongly favors 3’-end formation. Nevertheless, our results together with emerging evidence suggest that chromatin structure and transcription termination are mechanistically coupled. Our observation that mutations within the KOW-stalk surface produce defects in both processes supports this model.

This work has identified a potentially novel function for the Pol II stalk in chromatin regulation through characterization of alleles that result in cryptic initiation. The identification of novel alleles in *spt5* that also result in cryptic initiation near the Pol II stalk, as well as enrichment of Spt16, Nap1 and histone proteins in KOW pulldowns, supports a model in which this surface is involved in regulating chromatin structure during transcription. One prominent feature of histone interacting proteins is the presence of acidic regions that promote interactions with the basic histone proteins. The surface composed of Rpb4/7 and KOW2/3 is predominantly acidic in nature according to a charge distribution map (Supplemental Figure 3), and the surface-exposed residues altered in our cryptic initiation assay are both acidic (Rpb7-D166, Spt5-E546), suggesting this surface has the potential to engage histones. Interestingly, a recent structural model of isolated native Pol II ECs shows Rpb7 directly contacting an upstream nucleosome (Kujirai et al. 2025). In this structure, Rpb7-D166 lies nearby, but not in direct contact with the incoming nucleosome. Spt5 and other elongation factors were not resolved in this structure, and the authors interpreted this to be evidence of elongation factor-independent nucleosome reassembly via template looping. The authors note, however, that elongation factors may have dissociated during sample preparation. An alternative model for the cryptic initiation phenotype we observed is that KOW2-3 and Rpb4/7 do not directly facilitate nucleosome passage themselves but form a platform for the assembly of other chromatin modifying factors, thereby exerting an indirect effect on chromatin structure. Supporting this model, our mass spectrometry data identified the histone chaperones Nap1 and Spt16 as enriched in the KOW2-3 pulldown. Further, recent structural evidence shows Spt6’s cognate binding partner Iws1/Spn1 binding to Spt5’s NGN and KOW2 domains simultaneously with Spt6’s N-terminal helices (Ehara et al. 2022). Additionally, Spt6 has been shown through structural studies to make extensive contacts with Rpb4/7 (Ehara et al. 2017; Vos et al. 2018). Notably, these structures show contacts between Spt6 and Rpb7-D166, as well as Spn1 and Spt5-E546, suggesting a potential mechanism for chromatin disruption in *rpb7-D166G* and *spt5-E546K.* Future experiments should distinguish direct from indirect roles of Spt5 and Rpb7 in nucleosome traversal and reassembly, for example using single-molecule optical tweezers assays (Chen et al. 2019; Burgos-Bravo et al. 2025).

Taken together, these results support a model in which the surface formed by Spt5 KOW2-3 and the Pol II stalk regulates chromatin integrity and termination in a context-dependent manner. Future studies should consider that this structure acts to provoke interactions with tertiary factors to execute these functions. This model does not preclude, however, direct effects on chromatin or RNA, either via direct interactions with nucleosomes through its acidic surface, or direct contacts with RNA to regulate termination outcome, as the stalk and Spt5 have both been shown to bind RNA (Meka et al. 2005; Ujvári and Luse 2006; Blythe et al. 2016; Zuber et al. 2018). Future work should aim to deconvolute direct vs indirect effects of this surface on co-transcriptional processes and define the contexts in which these mechanisms are regulated.

## Data Availability Statement

All Strains and plasmids are available upon request. The authors affirm that all data necessary for confirming the conclusions of the article are present within the article, figures, and tables. A detailed list of yeast strains used throughout this manuscript is available in Supplementary Table 1. A detailed list of plasmids used is available in Supplementary Table 2. A list of all oligonucleotides used in this study is available in Supplementary Table 3. A complete list of proteins identified in MudPIT mass spectrometry experiments are available as Supplementary Files 1 (K2K3), 2 (L2K4), and 3 (BSA).

## Acknowledgements

We would like to acknowledge Dr. Craig Kaplan for providing the *RPB7* strains used in this study, Dr. Jinhua Fu for providing expression constructs for the affinity chromatography experiments, and Dr. David Brow for providing the Nrd1 antibody and *NRD1* plasmid used in this work.

## Study Funding

This work was supported by internal UCSC funds (GAH), NIH grants 5R01GM032543 (CJB), NIH P41 GM103533 (JYIII) and the Howard Hughes Medical Institute (CJB).

## Conflict of Interests

The authors declare no conflict of interests in relation to the work described here.

---

**Supplementary Figure 1:** Multiple-sequence alignment of the sequence spanning Rpb7 amino acids 80-171, which includes G149D, D166G, L168S, and Spt5 amino acids 495-638, including E546K, G587D. Red highlight indicates universal conservation among organisms analyzed. Blue highlight indicates residues that are surface exposed, red highlight indicates residues that lie at the interface of KOW3 and Rpb7. Alignment was generated via Uniprot (UniProt: the Universal Protein Knowledgebase in 2025 | Nucleic Acids Research | Oxford Academic).

**Supplementary Figure 2:** Silver-stained gradient gels containing the input, washes, and KCl gradient elutions of Spt5-KOW affinity chromatography and BSA control.

**Supplementary Figure 3:** Charge distribution map of Rpb4, Rpb7 and Spt5 KOW2-3. Red indicates negative charge; blue indicates positive charge. Circles indicate locations of amino acid changes in *spt5, rpb7* mutants. PDB: 5OIK (Bernecky et al. 2017).

## References

Allepuz-Fuster P, O’Brien MJ, González-Polo N, Pereira B, Dhoondia Z, Ansari A, Calvo O. 2019. RNA polymerase II plays an active role in the formation of gene loops through the Rpb4 subunit. Nucleic Acids Res. 47(17):8975–8987. doi:10.1093/nar/gkz597.

Armache K-J, Mitterweger S, Meinhart A, Cramer P. 2005. Structures of complete RNA polymerase II and its subcomplex, Rpb4/7. J Biol Chem. 280(8):7131–7134. doi:10.1074/jbc.M413038200.

Bernecky C, Plitzko JM, Cramer P. 2017. Structure of a transcribing RNA polymerase II-DSIF complex reveals a multidentate DNA-RNA clamp. Nat Struct Mol Biol. 24(10):809–815. doi:10.1038/nsmb.3465.

Blythe AJ, Yazar-Klosinski B, Webster MW, Chen E, Vandevenne M, Bendak K, Mackay JP, Hartzog GA, Vrielink A. 2016. The yeast transcription elongation factor Spt4/5 is a sequence-specific RNA binding protein. Protein Sci. 25(9):1710–1721. doi:10.1002/pro.2976.

Braberg H, Jin H, Moehle EA, Chan YA, Wang S, Shales M, Benschop JJ, Morris JH, Qiu C, Hu F, et al. 2013. From Structure to Systems: High-Resolution, Quantitative Genetic Analysis of RNA Polymerase II. Cell. 154(4):775–788. doi:10.1016/j.cell.2013.07.033.

Burgos-Bravo F, Tong AB, Li C, Díaz-Celis C, Kaplan CD, LeRoy G, Reinberg D, Bustamante C. 2025. FACT weakens the nucleosomal barrier to transcription and preserves its integrity by forming a hexasome-like intermediate. Mol Cell. 85(11):2097–2109.e8. doi:10.1016/j.molcel.2025.05.002.

Calvo O. 2020. RNA polymerase II phosphorylation and gene looping: new roles for the Rpb4/7 heterodimer in regulating gene expression. Curr Genet. 66(5):927–937. doi:10.1007/s00294-020-01084-w.

Chen Z, Gabizon R, Brown AI, Lee A, Song A, Díaz-Celis C, Kaplan CD, Koslover EF, Yao T, Bustamante C. 2019. High-resolution and high-accuracy topographic and transcriptional maps of the nucleosome barrier. eLife. 8:e48281. doi:10.7554/eLife.48281.

Cheung V, Chua G, Batada NN, Landry CR, Michnick SW, Hughes TR, Winston F. 2008. Chromatin- and Transcription-Related Factors Repress Transcription from within Coding Regions throughout the Saccharomyces cerevisiae Genome. PLOS Biology. 6(11):e277. doi:10.1371/journal.pbio.0060277.

Choder M. 2004. Rpb4 and Rpb7: subunits of RNA polymerase II and beyond. Trends in Biochemical Sciences. 29(12):674–681. doi:10.1016/j.tibs.2004.10.007.

Choder M, Young RA. 1993. A portion of RNA polymerase II molecules has a component essential for stress responses and stress survival. Mol Cell Biol. 13(11):6984–6991. doi:10.1128/mcb.13.11.6984-6991.1993.

Crickard JB, Lee J, Lee T-H, Reese JC. 2017. The elongation factor Spt4/5 regulates RNA polymerase II transcription through the nucleosome. Nucleic Acids Research. 45(11):6362–6374. doi:10.1093/nar/gkx220.

Cui Y, Denis CL. 2003. In Vivo Evidence that Defects in the Transcriptional Elongation Factors RPB2, TFIIS, and SPT5 Enhance Upstream Poly(A) Site Utilization. Mol Cell Biol. 23(21):7887–7901. doi:10.1128/MCB.23.21.7887-7901.2003.

Ding B, LeJeune D, Li S. 2010. The C-terminal repeat domain of Spt5 plays an important role in suppression of Rad26-independent transcription coupled repair. J Biol Chem. 285(8):5317–5326. doi:10.1074/jbc.M109.082818.

Edwards AM, Kane CM, Young RA, Kornberg RD. 1991. Two dissociable subunits of yeast RNA polymerase II stimulate the initiation of transcription at a promoter in vitro. J Biol Chem. 266(1):71–75.

Ehara H, Kujirai T, Shirouzu M, Kurumizaka H, Sekine S-I. 2022. Structural basis of nucleosome disassembly and reassembly by RNAPII elongation complex with FACT. Science. 377(6611):eabp9466. doi:10.1126/science.abp9466.

Ehara H, Yokoyama T, Shigematsu H, Yokoyama S, Shirouzu M, Sekine S-I. 2017. Structure of the complete elongation complex of RNA polymerase II with basal factors. Science. 357(6354):921–924. doi:10.1126/science.aan8552.

Evrin C, Serra-Cardona A, Duan S, Mukherjee PP, Zhang Z, Labib KPM. 2022. Spt5 histone binding activity preserves chromatin during transcription by RNA polymerase II. EMBO J. 41(5):e109783. doi:10.15252/embj.2021109783.

Geisberg JV, Moqtaderi Z, Fong N, Erickson B, Bentley DL, Struhl K. 2022. Nucleotide-level linkage of transcriptional elongation and polyadenylation. Elife. 11:e83153. doi:10.7554/eLife.83153.

Geisberg JV, Moqtaderi Z, Struhl K. 2020. The transcriptional elongation rate regulates alternative polyadenylation in yeast. Elife. 9:e59810. doi:10.7554/eLife.59810.

Geisberg JV, Moqtaderi Z, Struhl K. 2024. Chromatin regulates alternative polyadenylation via the RNA polymerase II elongation rate. Proc Natl Acad Sci U S A. 121(21):e2405827121. doi:10.1073/pnas.2405827121.

Gong W, Li S. 2023. Rpb7 represses transcription-coupled nucleotide excision repair. J Biol Chem. 299(8):104969. doi:10.1016/j.jbc.2023.104969.

Greger IH, Proudfoot NJ. 1998. Poly(A) signals control both transcriptional termination and initiation between the tandem GAL10 and GAL7 genes of Saccharomyces cerevisiae. EMBO J. 17(16):4771–4779. doi:10.1093/emboj/17.16.4771.

Hartzog GA, Fu J. 2013. The Spt4-Spt5 complex: a multi-faceted regulator of transcription elongation. Biochim Biophys Acta. 1829(1):105–115. doi:10.1016/j.bbagrm.2012.08.007.

Heo D, Yoo I, Kong J, Lidschreiber M, Mayer A, Choi B-Y, Hahn Y, Cramer P, Buratowski S, Kim M. 2013. The RNA polymerase II C-terminal domain-interacting domain of yeast Nrd1 contributes to the choice of termination pathway and couples to RNA processing by the nuclear exosome. J Biol Chem. 288(51):36676–36690. doi:10.1074/jbc.M113.508267.

Kaplan CD, Holland MJ, Winston F. 2005. Interaction between Transcription Elongation Factors and mRNA 3′-End Formation at the Saccharomyces cerevisiae GAL10-GAL7 Locus *. Journal of Biological Chemistry. 280(2):913–922. doi:10.1074/jbc.M411108200.

Kolodziej PA, Woychik N, Liao SM, Young RA. 1990. RNA polymerase II subunit composition, stoichiometry, and phosphorylation. Mol Cell Biol. 10(5):1915–1920. doi:10.1128/mcb.10.5.1915-1920.1990.

Krogan NJ, Kim M, Ahn SH, Zhong G, Kobor MS, Cagney G, Emili A, Shilatifard A, Buratowski S, Greenblatt JF. 2002. RNA Polymerase II Elongation Factors of Saccharomyces cerevisiae: a Targeted Proteomics Approach. Mol Cell Biol. 22(20):6979–6992. doi:10.1128/MCB.22.20.6979-6992.2002.

Kujirai T, Kato J, Yamamoto K, Hirai S, Fujii T, Maehara K, Harada A, Negishi L, Ogasawara M, Yamaguchi Y, et al. 2025. Multiple structures of RNA polymerase II isolated from human nuclei by ChIP-CryoEM analysis. Nat Commun. 16(1):4724. doi:10.1038/s41467-025-59580-x.

Lasko P. 2010. Tudor Domain. Current Biology. 20(16):R666–R667. doi:10.1016/j.cub.2010.05.056.

Lee KY, Chopra A, Burke GL, Chen Z, Greenblatt JF, Biggar KK, Meneghini MD. 2020. A crucial RNA-binding lysine residue in the Nab3 RRM domain undergoes SET1 and SET3-responsive methylation. Nucleic Acids Research. 48(6):2897–2911. doi:10.1093/nar/gkaa029.

Leporé N, Lafontaine DLJ. 2011. A Functional Interface at the rDNA Connects rRNA Synthesis, Pre-rRNA Processing and Nucleolar Surveillance in Budding Yeast. PLOS ONE. 6(9):e24962. doi:10.1371/journal.pone.0024962.

Li W, Giles C, Li S. 2014. Insights into how Spt5 functions in transcription elongation and repressing transcription coupled DNA repair. Nucleic Acids Res. 42(11):7069–7083. doi:10.1093/nar/gku333.

Lindstrom DL, Squazzo SL, Muster N, Burckin TA, Wachter KC, Emigh CA, McCleery JA, Yates JR, Hartzog GA. 2003. Dual Roles for Spt5 in Pre-mRNA Processing and Transcription Elongation Revealed by Identification of Spt5-Associated Proteins. Mol Cell Biol. 23(4):1368–1378. doi:10.1128/MCB.23.4.1368-1378.2003.

Malone EA, Fassler JS, Winston F. 1993. Molecular and genetic characterization of SPT4, a gene important for transcription initiation in Saccharomyces cerevisiae. Mol Gen Genet. 237(3):449–459. doi:10.1007/BF00279450.

Mayer A, Schreieck, Amelie, Lidschreiber, Michael, Leike, Kristin, Martin, Dietmar E., and Cramer P. 2012. The Spt5 C-Terminal Region Recruits Yeast 3′ RNA Cleavage Factor I. Molecular and Cellular Biology. 32(7):1321–1331. doi:10.1128/MCB.06310-11.

Meka H, Werner F, Cordell SC, Onesti S, Brick P. 2005. Crystal structure and RNA binding of the Rpb4/Rpb7 subunits of human RNA polymerase II. Nucleic Acids Res. 33(19):6435–6444. doi:10.1093/nar/gki945.

Meyer PA, Li S, Zhang M, Yamada K, Takagi Y, Hartzog GA, Fu J. 2015. Structures and Functions of the Multiple KOW Domains of Transcription Elongation Factor Spt5. Mol Cell Biol. 35(19):3354–3369. doi:10.1128/MCB.00520-15.

Mitsuzawa H, Kanda E, Ishihama A. 2003. Rpb7 subunit of RNA polymerase II interacts with an RNA-binding protein involved in processing of transcripts. Nucleic Acids Res. 31(16):4696–4701.

Moqtaderi Z, Geisberg JV, Struhl K. 2026. Genetic analysis of polyadenylation patterns reveals distinct classes of yeast genes and local chromatin effects on Pol II elongation. Genetics.:iyag057. doi:10.1093/genetics/iyag057.

Mosley AL, Hunter GO, Sardiu ME, Smolle M, Workman JL, Florens L, Washburn MP. 2013. Quantitative Proteomics Demonstrates That the RNA Polymerase II Subunits Rpb4 and Rpb7 Dissociate during Transcriptional Elongation. Mol Cell Proteomics. 12(6):1530–1538. doi:10.1074/mcp.M112.024034.

Pandey V, Punniyamoorthy S, Pokharel YR. 2023. Emerging Roles of SPT5 in Transcription. Cell Physiol Biochem. 57(5):395–408. doi:10.33594/000000665.

Qiu Y, Gilmour DS. 2017. Identification of Regions in the Spt5 Subunit of DRB Sensitivity-inducing Factor (DSIF) That Are Involved in Promoter-proximal Pausing. J Biol Chem. 292(13):5555–5570. doi:10.1074/jbc.M116.760751.

Rosen GA, Baek I, Friedman LJ, Joo YJ, Buratowski S, Gelles J. 2020. Dynamics of RNA polymerase II and elongation factor Spt4/5 recruitment during activator-dependent transcription. Proc Natl Acad Sci U S A. 117(51):32348–32357. doi:10.1073/pnas.2011224117.

Runner VM, Podolny V, Buratowski S. 2008. The Rpb4 Subunit of RNA Polymerase II Contributes to Cotranscriptional Recruitment of 3′ Processing Factors. Mol Cell Biol. 28(6):1883–1891. doi:10.1128/MCB.01714-07.

Schier AC, Taatjes DJ. 2020. Structure and mechanism of the RNA polymerase II transcription machinery. Genes Dev. 34(7–8):465–488. doi:10.1101/gad.335679.119.

Sharma N, Kumari R. 2013. Rpb4 and Rpb7: multifunctional subunits of RNA polymerase II. Critical Reviews in Microbiology. 39(4):362–372. doi:10.3109/1040841x.2012.711742.

Singh P, Chaudhuri A, Banerjea M, Marathe N, Das B. 2021. Nrd1p identifies aberrant and natural exosomal target messages during the nuclear mRNA surveillance in Saccharomyces cerevisiae. Nucleic Acids Res. 49(20):11512–11536. doi:10.1093/nar/gkab930.

Song A, Chen FX. 2022. The pleiotropic roles of SPT5 in transcription. Transcription. 13(1–3):53–69. doi:10.1080/21541264.2022.2103366.

Steinmetz EJ, Brow DA. 1996. Repression of Gene Expression by an Exogenous Sequence Element Acting in Concert with a Heterogeneous Nuclear Ribonucleoprotein-Like Protein, Nrd1, and the Putative Helicase Sen1. Molecular and Cellular Biology. 16(12):6993–7003. doi:10.1128/MCB.16.12.6993.

Steinmetz EJ, Brow DA. 1998. Control of pre-mRNA accumulation by the essential yeast protein Nrd1 requires high-affinity transcript binding and a domain implicated in RNA polymerase II association. Proceedings of the National Academy of Sciences. 95(12):6699–6704. doi:10.1073/pnas.95.12.6699.

Steinmetz EJ, Conrad NK, Brow DA, Corden JL. 2001. RNA-binding protein Nrd1 directs poly(A)-independent 3’-end formation of RNA polymerase II transcripts. Nature. 413(6853):327–331. doi:10.1038/35095090.

Tan Q, Prysak MH, Woychik NA. 2003. Loss of the Rpb4/Rpb7 subcomplex in a mutant form of the Rpb6 subunit shared by RNA polymerases I, II, and III. Mol Cell Biol. 23(9):3329–3338. doi:10.1128/MCB.23.9.3329-3338.2003.

Ujvári A, Luse DS. 2006. RNA emerging from the active site of RNA polymerase II interacts with the Rpb7 subunit. Nat Struct Mol Biol. 13(1):49–54. doi:10.1038/nsmb1026.

UniProt: the Universal Protein Knowledgebase in 2025 | Nucleic Acids Research | Oxford Academic. [accessed 2026 Mar 8]. https://academic.oup.com/nar/article/53/D1/D609/7902999?login=false.

Vasiljeva L, Buratowski S. 2006. Nrd1 interacts with the nuclear exosome for 3’ processing of RNA polymerase II transcripts. Mol Cell. 21(2):239–248. doi:10.1016/j.molcel.2005.11.028.

Venette-Smith NL, Vishwakarma RK, Venkatakrishnan V, Dollinger R, Schultz J, Babitzke P, Anand G, Gilmour DS, Armache J-P, Murakami KS. 2025. Structural Characterization of Native RNA Polymerase II Transcription Complexes and Nucleosomes in Drosophila melanogaster.: 2025.02.03.636274. doi:10.1101/2025.02.03.636274. [accessed 2025 June 2]. https://www.biorxiv.org/content/10.1101/2025.02.03.636274v2.

Vos SM, Farnung L, Boehning M, Wigge C, Linden A, Urlaub H, Cramer P. 2018. Structure of activated transcription complex Pol II–DSIF–PAF–SPT6. Nature. 560(7720):607–612. doi:10.1038/s41586-018-0440-4.

Wang B, Artsimovitch I. 2020. NusG, an Ancient Yet Rapidly Evolving Transcription Factor. Front Microbiol. 11:619618. doi:10.3389/fmicb.2020.619618.

Zeng Y, Zhang H-W, Wu X-X, Zhang Y. 2024. Structural basis of exoribonuclease-mediated mRNA transcription termination. Nature. 628(8009):887–893. doi:10.1038/s41586-024-07240-3.

Zuber PK, Hahn L, Reinl A, Schweimer K, Knauer SH, Gottesman ME, Rösch P, Wöhrl BM. 2018. Structure and nucleic acid binding properties of KOW domains 4 and 6–7 of human transcription elongation factor DSIF. Sci Rep. 8(1):11660. doi:10.1038/s41598-018-30042-3.

